# Longitudinal brain structure changes in Parkinson’s disease: a replication study

**DOI:** 10.1101/2023.04.28.538743

**Authors:** Andrzej Sokołowski, Nikhil Bhagwat, Yohan Chatelain, Mathieu Dugré, Alexandru Hanganu, Oury Monchi, Brent McPherson, Michelle Wang, Jean-Baptiste Poline, Madeleine Sharp, Tristan Glatard

## Abstract

**Context:** An existing major challenge in Parkinson’s disease (PD) research is the identification of biomarkers of disease progression. While Magnetic Resonance Imaging (MRI) is a potential source of PD biomarkers, none of the MRI measures of PD are robust enough to warrant their adoption in clinical research. This study is part of a project that aims to replicate 11 PD studies reviewed in a recent survey (*JAMA neurology, 78*(10) 2021) to investigate the robustness of PD neuroimaging findings to data and analytical variations.

**Objective:** This study attempts to replicate the results in Hanganu et al. (*Brain, 137*(4) 2014) using data from the Parkinson’s Progression Markers Initiative (PPMI).

**Methods:** Using 25 PD subjects and 18 healthy controls, we analyzed the rate of change of cortical thickness and of the volume of subcortical structures, and we measured the relationship between MRI structural changes and cognitive decline. We compared our findings to the results in the original study.

**Results:** Similarly to the original study, PD patients with mild cognitive impairment (MCI) exhibited increased cortical thinning over time compared to patients without MCI in the right middle temporal gyrus, insula, and precuneus. (2) The rate of cortical thinning in the left inferior temporal and precentral gyri in PD patients correlated with the change in cognitive performance. (3) There were no group differences in the change of subcortical volumes. (4) We did not find a relationship between the change in subcortical volumes and the change in cognitive performance.

**Conclusion:** Despite important differences in the dataset used in this replication study, and despite differences in sample size, we were able to partially replicate the original results. We produced a publicly available reproducible notebook allowing researchers to further investigate the reproducibility of the results in Hanganu et al. (2014) when more data becomes available in PPMI.

## Introduction

Parkinson’s disease (PD), one of the most common neurodegenerative diseases, is often characterized by akinesia, bradykinesia, tremor, and is commonly associated with mild cognitive impairment which significantly decrease overall quality of life [1]. These symptoms are accompanied by atrophy of the cortical and subcortical brain structures as well as cortical thinning [2]. As a result, there has been interest in determining whether MRI measures of atrophy can be used as a biomarker of cognitive decline. Overall, gray matter atrophy and cortical thinning are present in early PD, while frontal atrophy and temporoparietal thinning are associated with cognitive impairment in PD [3].

MRI-derived measures of the structural brain changes occuring in PD have emerged as potential diagnostic and prognostic tools to understand the trajectory of PD. Structural imaging, especially regional cortical thickness and loss of gray matter volume, has been considered helpful in determining a PD diagnosis, progression prognosis, and distinguishing PD from other dementias [2]. However, the need to further investigate the sensitivity, reliability, effect of confounding factors, and overall generalizability of these progression measures has been highlighted (e.g., [3]) and such rigorous validation is likely a factor preventing the adoption of MRI measures as outcome measures in PD clinical research.

The replication of neuroimaging findings has been challenged in multiple ways in recent years. For example, in the study of Botvinik-Nezer et al. [4], 70 independent teams were asked to analyze the same dataset using the methods of their choice. Results obtained across research teams did not concur on five out of the nine ex-ante hypotheses, reaching agreement levels ranging from 21% to 37%. Furthermore, the identification of regional brain atrophy in PD has been of interest as a possible marker of certain symptoms of PD and of the progression of PD [2]. However, studies conducted in non-PD populations have shown that estimates of regional volume [5,6] and of cortical thickness vary depending on the software toolbox [7,8]. Overall, a range of factors matter in the replicability of neuroimaging findings, including computational environments [9,10], analysis tools and versions [11,12], statistical models [13], and study populations [14].

This study is a part of a reproducibility evaluation project that aims to replicate 11 structural MRI measures of PD reviewed in Mitchel et al. [2]. The goal of the present study is to replicate the work by Hanganu et al. [15] to test whether prior findings regarding structural MRI-derived PD biomarkers replicate in a different dataset using similar analytical methods. Hanganu et al. [15] compared the change of gray matter volume and cortical thinning over time between PD patients with mild cognitive impairment (PD-MCI), PD patients without mild cognitive impairment (PD-non-MCI), and healthy older controls (HC); and also tested the relationship between longitudinal structural changes and cognitive decline in the PD patients. They reported four main findings: (Finding 1) an increased rate of cortical thinning in PD patients with mild cognitive impairment compared to PD patients without MCI (mainly affecting the right temporal regions, insula, and inferior frontal gyrus), and compared to healthy controls (mainly in the right temporal regions and supplementary motor area); (Finding 2) a correlation between the change in Montreal Cognitive Assessment (MoCA) scores and cortical thinning in the bilateral temporal lobe, right occipital medial lobe, and the left postcentral gyrus in PD patients; (Finding 3) an increased loss of the amygdala and nucleus accumbens volumes as well as overall cortical thickness for the PD-MCI group compared to HC; (Finding 4) a correlation between the change of the right amygdala and thalamus volumes and the change in MoCA scores in PD patients.

The results of Hanganu et al. [15] are of clinical interest because they provide insight into the relationship between structural brain changes and cognitive impairment, thus highlighting possible neural substrates of PD-related cognitive impairment [16]. Our study addresses the issue of MRI measure replicability, investigates crucial elements of reporting the study to make it replicable, and discusses the impact of study design decisions on the replicability. The goal of our study was to attempt to replicate the original findings using a different cohort. We used open data from the Parkinson’s Progression Markers Initiative (PPMI; www.ppmi-info.org) in order to construct a similar patient cohort as that used in the original study and we followed the data processing methods and statistical analyses from the original study.

## Methods

### Participants

The original study included 15 PD-non-MCI, 17 PD-MCI and 18 HC. In order to reconstruct this cohort, PD patients and HC were selected from PPMI to attempt to match the sample size and demographics of the groups in the original study. The following criteria were used to define the PD cohorts: clinical diagnosis of PD, available T1-weighted images at two research visits, Hoehn and Yahr stage I and II (the stage was stable across the two visits for each patient), testing performed at PD OFF state, available MoCA scores, and the absence of any other neurological condition. Data was collected after approval of the local ethics committees of the PPMI’s participating sites. All participants provided written informed consent. This study was conducted in accordance with the Declaration of Helsinki and was exempt from the Concordia University’s Research Ethics Unit.

Patients were divided into PD-MCI and PD-non-MCI groups. In the original study, MCI was diagnosed on the basis of the presence of subjective complaints of cognitive impairment, objective impairment on two or more neuropsychological tests in one domain of cognitive function and the absence of dementia. In the PPMI dataset, patients are already classified as having MCI or not using a very similar criteria for classification, and thus the existing classification was used. Diagnosis of MCI in the PPMI is determined based on the following criteria: impairment in at least one cognitive domain, decline from pre-morbid function, and lack of significant impact of cognitive impairment on daily function.

Ten PD-MCI (*M* age = 67.6; *SD* = 5.8), 15 PD-non-MCI (*M* age = 63.4; *SD* = 9.4), and 18 HC participants were selected (*M* age = 66.9; *SD* = 6.1). PD-non-MCI and HC group sample sizes match those of the original study but an insufficient number of PD-MCI patients were identified in the PPMI dataset that met all the original inclusion criteria, thus our sample is smaller than the original sample (n=10 vs n=17). Descriptive statistics are reported in Table 1.

**Table 1.**
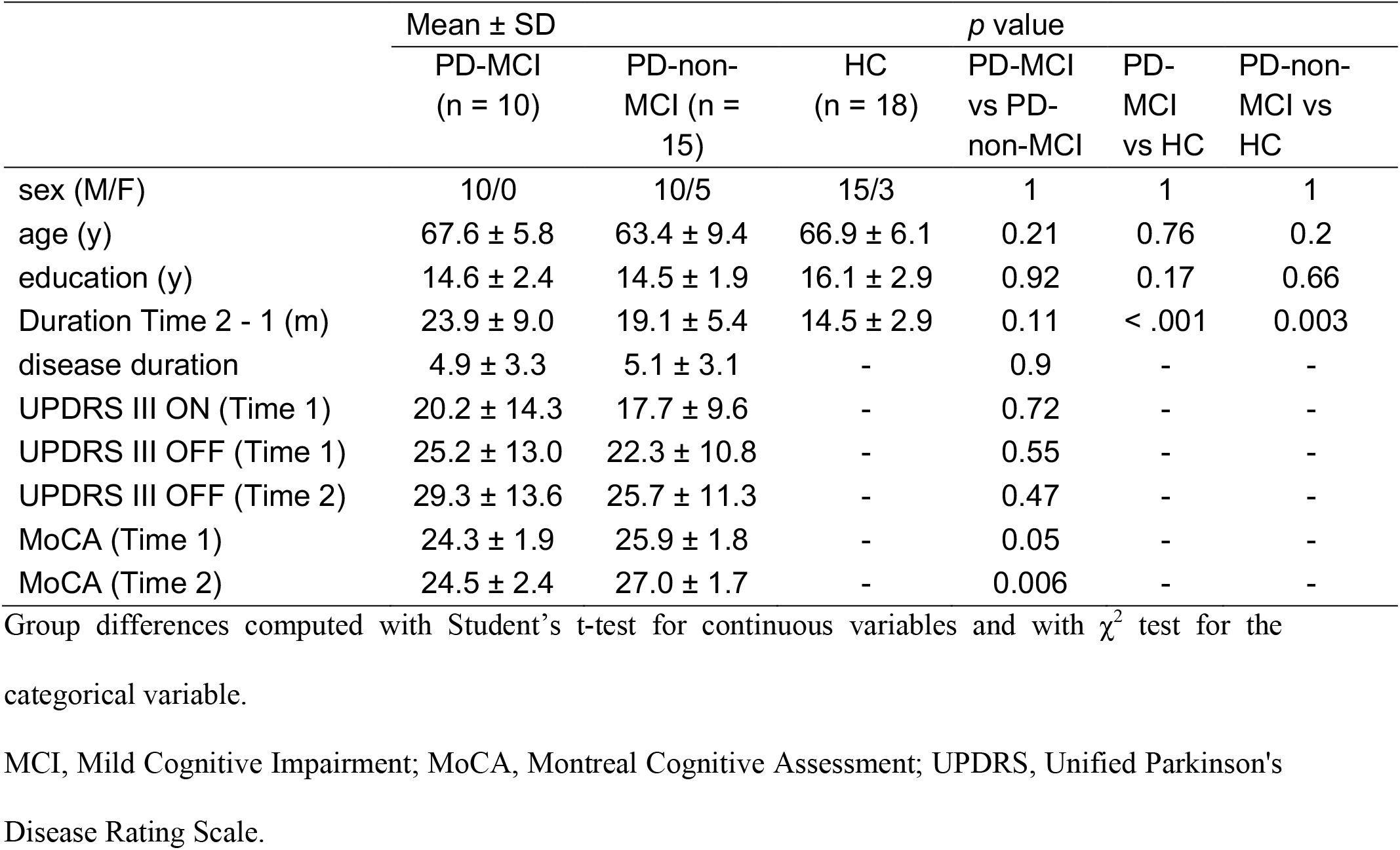
Descriptive statistics

### Image acquisition and preprocessing

MRI images were taken from the PPMI which uses a standardized study protocol and the following parameters: repetition time = 2.3 s, echo time = 2.98 s, inversion time = 0.9 s, slice thickness = 1 mm, number of slices = 192, field of view = 256 mm, and matrix size = 256 × 256. However, since PPMI is a multisite project there may be slight differences in the sites’ setup. Scans were acquired using different 3T scanners (Philips Achieva n=2; Siemens Prisma fit n=8; Siemens Prisma n=4; Siemens Skyra n=2; Siemens TrioTim n=64; Siemens Verio n=6). There were scans with echo time (TE) that diverged from the standardized protocol: one image with TE = 2.52 s, one image with TE = 1.91 s, three images with TE = 2.91 s, one image with TE = 2.93 s, four images with TE = 2.95 s, four images with TE = 2.96 s, two images with TE = 3.06s. Additionally, two images had TE = 2.94 s, TR = 6.49 s, and TE = 2.91 s, TR 6.26 s.

T1-weighted brain images were processed using FreeSurfer 7.1.1 [17]. The longitudinal preprocessing stream was used to calculate the change in cortical thinning and subcortical volumes [18]. FreeSurfer’s recon-all function was used for cortical reconstruction. First, all timepoints were processed cross-sectionally with the default workflow, then an unbiased template from the two timepoints was created for each subject, finally data was processed longitudinally. Specifically an unbiased within-subject template space and image [19] is created using robust, inverse consistent registration [20]. Several processing steps, such as skull stripping, Talairach transforms, atlas registration as well as spherical surface maps and parcellations are then initialized with common information from the within-subject template, significantly increasing reliability and statistical power [18]. The rate of change of cortical thickness between the two timepoints was calculated for each subject. Cortical thickness was smoothed with a 10 mm FWHM kernel. The original study also reported manual correction of misclassified tissue types, which was not performed in our study since the protocol for it was insufficient to replicate.

### Statistical analyses

Structural brain images and Montreal Cognitive Assessment (MoCA) scores from the initial and the follow-up visits were analyzed consistently with the 4 main findings reported in the original study. (Finding 1) We tested vertex-wise differences in the change of cortical thickness between HC, PD-MCI, and PD-non-MCI groups with an ANCOVA model. (Finding 2) We tested the correlation between the change of cortical thickness and the change of MoCA scores in PD-MCI, PD-non-MCI, and PD-all (all PD patients) groups. The time between the two visits was added as a covariate in the general linear models. Cluster-wise p-value threshold was used at the *p* < .05 level. The rate of change of the cortical thickness was calculated with the formula: (Thickness at Time 1 – Thickness at Time 2) / (Time 2 – Time 1). Subcortical volumes were adjusted for the estimated total intracranial volume as well as the averages of the two time points using regression-based correction, in line with the original study. (Finding 3) We tested the differences in regional volume changes between the three groups using t-tests and (Finding 4) measured the correlations between the change in MoCA scores and change of the subcortical volumes and cortical thickness in each group using Pearson correlation.

### Code availability

We used publicly available software to facilitate reproducing our study. Pandas v. 1.5.2 was used to define the cohort from PPMI data files. FreeSurfer 7.1.1 was used for image preprocessing and vertex-wise analyses. We used a containerized version of FreeSurfer managed by Boutiques 0.5.25 (doi:10.5281/zenodo.3839009). The containerized FreeSurfer analyses were executed through the Slurm batch manager on the Narval cluster (https://docs.alliancecan.ca/wiki/Narval/en) hosted at Calcul Québec and part of Digital Research Alliance of Canada.

The code and results are publicly available at https://github.com/LivingPark-MRI/hanganu-etal-2014. Data used in the notebook were downloaded directly from the PPMI and cannot be shared publicly due to its Data Usage Agreements preventing republishing data. We developed a Python package (LivingPark utils, available at https://github.com/LivingPark-MRI/livingpark-utils) to download and manipulate PPMI data directly from the original PPMI database. As a result, our notebook can be re-executed by anyone with a PPMI account.

## Results

### Vertex-wise results

We found numerous group differences in the rate of change of cortical thickness. The results are reported in Table 2 and Fig. 1.

**Table 2.** Vertex-wise group differences in the rate of change of cortical thickness

| Group and region | Size<br>(mm<sup>2</sup>) | MNI X | MNI Y | MNI Z | Max | clusterwise<br>*p*-value |
| --- | --- | --- | --- | --- | --- | --- |
| **PD-MCI vs PD-non-MCI** |  |  |  |  |  |  |
| L precuneus | 901.62 | -15.4 | -45.2 | 65.8 | 2.8711 | .0004 |
| R precentral | 808.31 | 45 | -4.9 | 49.8 | -4.3574 | .0026 |
| R insula | 689.83 | 36.1 | -16.4 | -3.4 | -4.285 | .0066 |
| R MTG | 636.56 | 42.1 | 7.1 | -37.1 | -4.7243 | .0108 |
| **PD-MCI vs HC** |  |  |  |  |  |  |
| L SPL | 1596.42 | -12.2 | -55.4 | 59.9 | 3.4112 | .0002 |
| L SFG | 816.25 | -8.3 | 35.1 | 34 | 4.2357 | .0006 |
| R precentral | 619.04 | 15 | -12.2 | 61.8 | 2.4693 | .0144 |
| R supramarginal | 578.61 | 37.3 | -29.7 | 39.5 | 3.0369 | .0221 |
| R SPL | 563.81 | 29.5 | -48.4 | 42.7 | 3.9183 | .0256 |

| PD-non-MCI vs HC |  |  |  |  |  |  |
| --- | --- | --- | --- | --- | --- | --- |
| R SMG | 1010.34 | 35.7 | -30.6 | 39.6 | 4.4171 | .0002 |
| R precentral | 580.49 | 43.5 | -9.4 | 46.2 | 3.8495 | .0211 |
| R precuneus | 537.11 | 8.6 | -71.2 | 40 | 3.6102 | .0384 |

HC, healthy controls; Max, maximum $-\log_{10}(p \text{ value})$ in the cluster; MTG, middle temporal gyrus; PD-non-MCI, Parkinson's disease without mild cognitive impairment; PD-MCI, Parkinson's disease with mild cognitive impairment; SFG, superior frontal gyrus; SMG, supramarginal gyrus; SPL, superior parietal lobule.

**Fig. 1.**
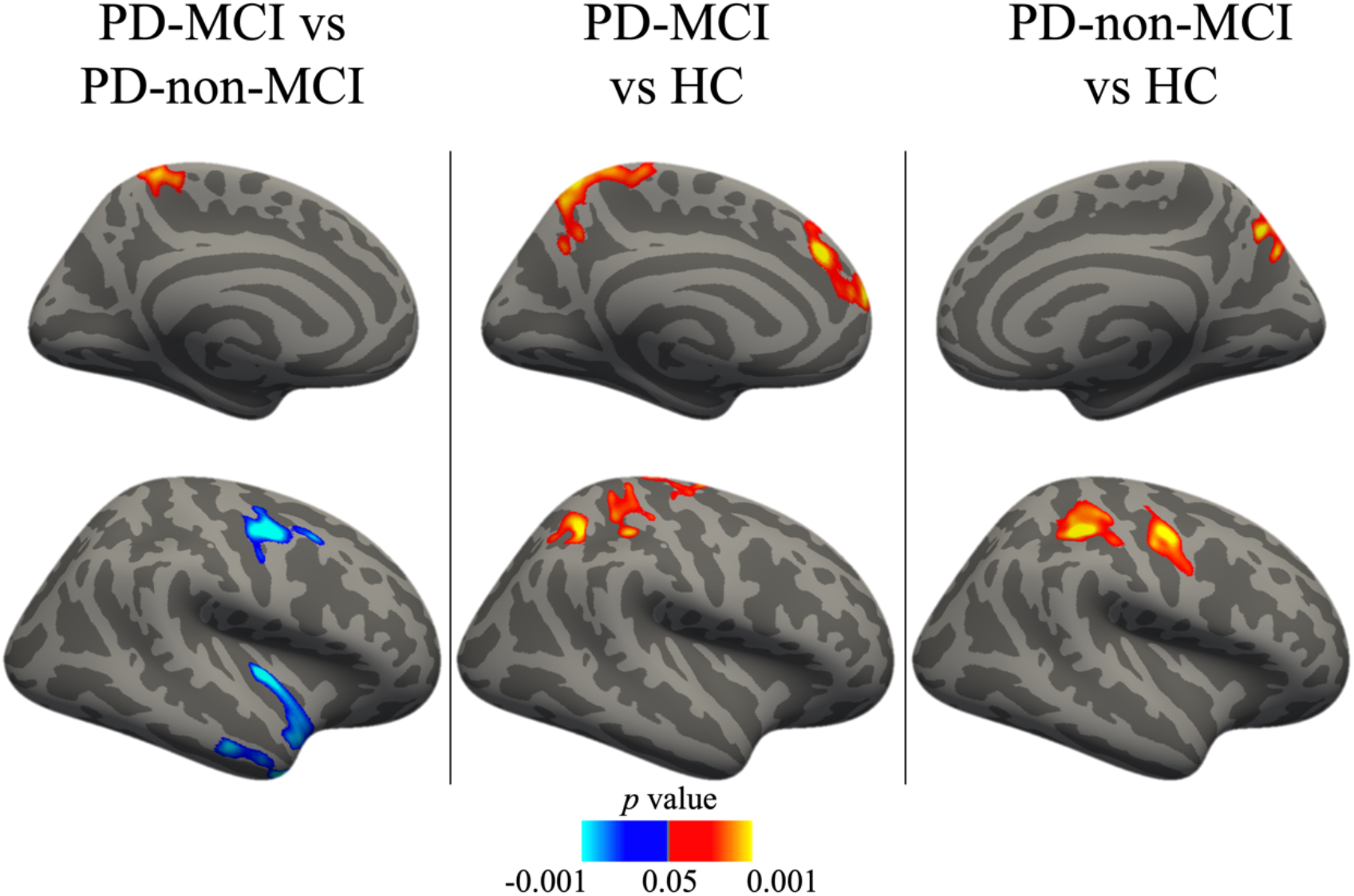
Vertex-wise group differences in the rate of change of cortical thickness. Cold colors represent increased cortical thinning in the first group compared to the second group, warm colors represent increased cortical thinning in the second group compared to the first group. Results corrected with the cluster-wise threshold (*p* < .05). HC, healthy controls; PD-MCI, Parkinson’s disease with mild cognitive impairment; PD-non-MCI, Parkinson’s disease without mild cognitive impairment.

#### PD-MCI vs. PD-non-MCI

The two patient groups differed in the rate of cortical thinning. The PD-MCI group, compared to PD-non-MCI patients, had an increased rate of cortical thinning in the right middle temporal and precentral gyri as well as right insula. PD-non-MCI group exhibited an increased thinning in the left precuneus compared to the PD-MCI group.

#### PD-MCI vs. HC

The HC group had increased cortical thinning in the right precentral and supramarginal gyri, left superior frontal gyrus, and bilateral superior parietal lobule, compared to the PD-MCI group.

#### PD-non-MCI vs. HC

The HC group had increased cortical thinning in the right precuneus as well as precentral and supramarginal gyri compared to the PD-non-MCI group.

There was a significant negative correlation between the change in MoCA scores and the rate of change of the right middle frontal gyrus thickness in the PD-MCI group (*p* < .05). There was a positive correlation between the MoCA and the rate of change of the left inferior temporal and precentral gyri across all PD patients (*p* < .05). The correlations were not significant in the PD-non-MCI group. The results are reported in Table 3 and Fig. 2.

**Table 3.** Vertex-wise correlation between the rate of change of cortical thickness and the change in Montreal Cognitive Assessment scores in PD patients

| Group and region | Size (mm<sup>2</sup>) | MNI X | MNI Y | MNI Z | Max | clusterwise $p$-value |
| --- | --- | --- | --- | --- | --- | --- |
| **PD-all** |  |  |  |  |  |  |
| L ITG | 592.67 | -53.3 | -20.1 | -33.1 | 3.4012 | .0177 |
| L precentral | 523.77 | -24.4 | -16.8 | 64.5 | 2.5887 | .0406 |
| **PD-MCI** |  |  |  |  |  |  |
| R rostral MTG | 749.67 | 30.1 | 43.1 | 16.9 | -4.0378 | .0004 |

ITG, inferior temporal gyrus; L, left; Max, maximum $-\log_{10}(p \text{ value})$ in the cluster; MCI, mild cognitive impairment; MTG, middle temporal gyrus; R, right.

**Fig. 2.**
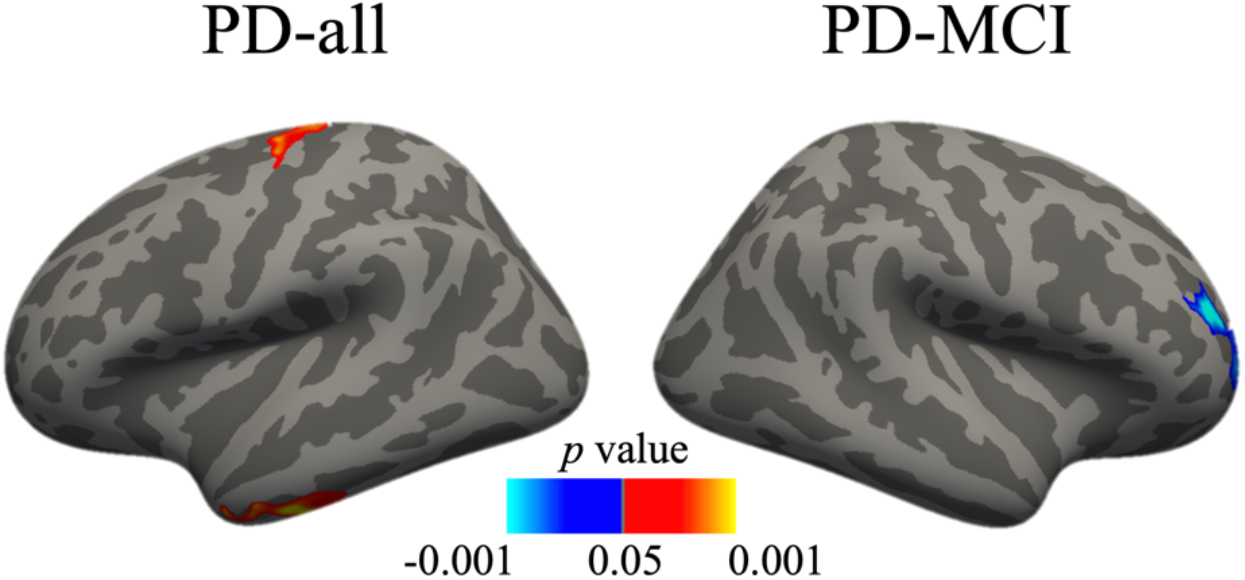
Vertex-wise correlation between the rate of change of cortical thickness and the change in Montreal Cognitive Assessment scores in the PD-all and PD-MCI groups.

### Volumetric results

Group comparisons of the volumetric change of subcortical regions revealed increased decrease of the thalamus volume in the PD-MCI group compared to PD-non-MCI (*p* = .01). There was a difference in the change of the putamen volume between the PD-non-MCI and HC groups (*p* = .02), with HC exhibiting increased thinning. The results are reported in Table 4.

**Table 4.** Group differences in the volumetric change of the subcortical regions and the overall cortical thickness

|  | PD-MCI |  | PD-non-MCI |  | HC |  | *p* values |  |  |
| --- | --- | --- | --- | --- | --- | --- | --- | --- | --- |
|  | mean | % | mean | % | mean | % | PD-MCI vs PD-non-MCI | PD-MCI vs HC | PD-non-MCI vs HC |
| CoTh | 0.006 | -0.57 | 0 | -0.11 | 0.006 | -0.21 | .44 | 1 | .5 |
| Thal | -223.24 | -2.56 | 210.34 | 1.25 | -51.26 | -0.68 | .01 | .28 | .09 |
| Caud | 36.33 | 0.51 | -12.99 | -0.25 | -9.36 | -0.2 | .63 | .56 | .95 |
| Put | -16.38 | -0.66 | 115.21 | 1.03 | -86.9 | -0.92 | .21 | .26 | .02 |
| Hipp | -50.43 | -1.76 | 4.56 | -1.14 | 24.22 | -0.49 | .43 | .22 | .74 |
| Amyg | -39.03 | -4.97 | -5.15 | -2.24 | 25.98 | -0.65 | .57 | .2 | .5 |
| Nacc | -9.89 | -1.23 | -3.06 | -0.82 | 8.04 | 0.23 | .81 | .52 | .58 |

Subcortical volumes are presented in mm<sup>3</sup>, cortical thickness in mm. Amyg, amygdala; Caud, caudate; CoTh, cortical thickness; Hipp, hippocampus; Nacc, nucleus accumbens; Put, putamen; Thal, thalamus.

Correlation analysis did not show any significant correlation between the change of MoCA scores and the change in volume of subcortical structures. Results are reported in Table 5.

**Table 5.** Correlation between the rate of change of the subcortical volumes and the change in Montreal Cognitive Assessment scores

| Structure | PD-all |  | PD-MCI |  | PD-non-MCI |  |
| --- | --- | --- | --- | --- | --- | --- |
|  | *r* | *p* | *r* | *p* | *r* | *p* |
| R Thalamus | -0.03 | .89 | 0.09 | .81 | 0.07 | .80 |
| R Caudate | 0.09 | .68 | -0.18 | .61 | 0.28 | .31 |
| R Putamen | 0.02 | .92 | 0.14 | .7 | 0.1 | .73 |
| R Pallidum | 0 | 1 | -0.07 | .84 | 0.33 | .22 |
| R Hippocampus | -0.1 | .64 | -0.04 | .92 | -0.05 | .85 |
| R Amygdala | -0.24 | .26 | -0.25 | .48 | -0.27 | .33 |
| R Nucleus accumbens | 0.04 | .87 | 0.38 | .27 | -0.12 | .66 |
| L Thalamus | 0.19 | .37 | 0.14 | .71 | 0.48 | .07 |
| L Caudate | 0 | 1 | -0.29 | .42 | 0.42 | .12 |
| L Putamen | -0.03 | .9 | -0.17 | .63 | 0.19 | .50 |
| L Pallidum | 0.05 | .81 | 0.02 | .97 | 0.15 | .61 |
| L Hippocampus | -0.01 | .96 | -0.08 | .82 | 0.07 | .81 |
| L Amygdala | 0.24 | .26 | 0.32 | .36 | 0.34 | .22 |
| L Nucleus accumbens | 0.02 | .92 | -0.07 | .85 | 0.12 | .67 |

L, left; MCI, mild cognitive impairment; PD-non-MCI, Parkinson's disease without mild cognitive impairment; PD-MCI, Parkinson's disease with mild cognitive impairment; R, right.

## Discussion

This study attempted to replicate the results of Hanganu et al. [15], which focused on the longitudinal changes in the cortical thickness and subcortical gray matter volume in PD patients. Group differences between PD patients with and without MCI were investigated along with the relationship between the structural changes and cognition. We used the analytic methods described in the paper and applied them to a different dataset of PD patients with and without MCI, and HC.

We have replicated the differences in the rate of cortical thinning between the PD-MCI and PD-non-MCI groups (Finding 1 in [15]), which supports the notion that these two patient groups differ in the rate of neurodegeneration. Patients with MCI had increased cortical thinning of the right MTG and insula, which overlaps with the regions reported in the original study. However, we did not replicate the group differences involving the HC group.

We found a relationship between the decrease in cognitive performance and cortical thinning of the left precentral gyrus and ITG in PD patients. The ITG cluster partially overlaps with the area reported in the original study. Although the structural changes were not found in the exact same brain voxels, the data supports the role of the thinning of the temporal regions in PD patients’ cognition (Finding 2 in [15]). Additionally, we found a relationship between the increase of the cognitive performance and cortical thinning of the right MFG in the PD-MCI group. This result was not reported in the original study and provides additional insight into potential structural changes in PD populations. Overall, we have replicated the original vertex-wise results (Findings 1 and 2 in [15]) to a certain extent.

Volumetric results (Findings 3 and 4 in [15]) were not replicated. The original study reported group differences in the change of overall cortical thickness as well as volumes of amygdala and nucleus accumbens. Instead, we found higher atrophy of thalamus in the PD-MCI compared to PD-non-MCI group, and higher putamen atrophy in the PD-MCI group compared to HC. Hanganu et al. [15] reported a correlation between the change in cognitive performance and the change in gray matter volume in the right thalamus and amygdala in PD patients. We found no significant correlations across all subcortical regions.

There are several factors that may explain the differences between the original results and the results of our replication. Most importantly, the two studies differ in the sample sizes for the PD-MCI group. We were unable to select enough PD patients with MCI that met all the inclusion criteria which reduced the chance to replicate the results. This was troublesome despite the fact that we used PPMI, one of the largest publicly available PD dataset. PPMI is a relatively new initiative and is lacking much longitudinal data. Our Jupyter notebook remains accessible and can be re-run as new data becomes available in PPMI. Importantly, current neuroscience recommends using larger sample sizes to avoid inflation of effect sizes. Sample sizes used in research are increasing and recent data suggest that brain-wide association studies may require as many as thousands of participants to define reliable brain-behavior relationships [21].

Differences between the samples may also affect replicability. There are no established clinical measures to infer disease severity in the brain, but disease duration, UPDRS score (a measure of symptom severity), and medication use are sometimes used as proxy measures. Average disease duration was very similar in our sample compared to the original study suggesting the patients were roughly matched. However, some patients had not yet started PD medications in our sample whereas all patients in the original study were already taking dopaminergic medications, and the UPDRS score was also a few points lower in our sample compared to the original study’s sample, both of which suggest that the replication sample we constructed from the PPMI cohort had slightly milder disease than the sample included in the original study. Our sample is also slightly older than the original cohort. Furthermore, despite using the same inclusion criteria as the original study, it is possible that other differences in sample characteristics may have contributed to the incomplete replication. Conclusions drawn from our data should not be expanded to different clinical populations (e.g., more severe PD patients).

Differences in neuroimaging data acquisition protocols and MRI scanners can also influence replicability. Data used in our study was acquired using different MRI scanners. Although PPMI uses a standardized protocol for data acquisition we cannot rule out the possibility that the reported differences may be related to the variability introduced by using various scanners, even though the vast majority of scans were acquired with a Siemens scanner. Ideally all participants should be scanned with the same machine but the benefits of collecting more data in a multisite project outweighs the advantages of a single-stage acquisition.

Software versions may also have impacted the results. FreeSurfer 5.3. for Centos 4 was used in the original study. This version of FreeSurfer was released 10 years ago and is no longer supported or recommended, hence we used version 7.1.1. for Centos 7 instead. Although the software version shouldn’t drastically change the clinical results, it is possible that it introduced variability during preprocessing or during the statistical analysis stage. We used the mri_glmfit-sim method to perform the analysis (including cluster correction) while the original study used the QDEC (Query, Design, Estimate, Contrast) method with mri_surfcluster. Our method is more stringent but also more reliable. It might have established more reliable borders between the gray and white matters. Filip et al. [22] reported structural group differences in data analyzed with FreeSurfer 5.3. which were not replicated with version 7.1.

Previous studies indicated that software version may impact structural brain analyses [7,9]. We will test the impact of software variability in our future work.

We thoroughly followed the original processing pipeline reported by Hanganu et al. [15]. Nevertheless, it is possible that we have missed some steps which could have influenced the results. We did not perform manual correction of misclassified brain tissue as this procedure cannot be objectively replicated. This may have impacted the analysis regarding smaller subcortical structures. There might be slight differences in the statistical models that we were not aware of. Any of the aforementioned discrepancies from the original study could have impacted the ability to replicate the results. Once the analyses were conducted, we contacted the authors of the original study to obtain their feedback, which importantly contributed to the discussion section.

Finally, there is a negative bias in replication studies, coming from the fact that researchers conducting replications focus on following original methods rather than getting positive results. Therefore, it is expected that more negative results are reported in replications than in original studies. We encourage scientists to follow all the steps and details from the original studies instead of simply aiming to replicate positive results.

We encountered multiple challenges while attempting to replicate the study by Hanganu et al. [15]. Analysis details are necessarily limited in the methods section of most papers which left us to infer some analytic steps. We also had difficulty constructing a similar cohort from publicly available PD data, which led to some differences in the patient characteristics between the original and replication samples, and to differences in sample size. We have published a Jupyter notebook that the research community can use to replicate our study. While respecting the PPMI data usage agreement, it clearly defines the criteria to define the study population (using the Pandas library), the preprocessing pipeline and software version (through containerized tools), and the statistical model that was used to obtain the results. Our notebook addresses some of the aforementioned challenges encountered during our study and can be re-run over time to update the study as more data gets added to PPMI.

We continue to investigate the replicability of MRI-derived PD biomarkers by replicating other clinical studies. Arafe et al. as well as Wang et al. report the results of their work in this Special Collection.

## Acknowledgment

This work was funded by the Michael J. Fox Foundation for Parkinson’s Research (MJFF-021134).

## Author Contributions

**Conceptualization**: Tristan Glatard, Jean-Baptiste Poline, Madeleine Sharp

**Data Curation**: Andrzej Sokołowski

**Formal Analysi**s: Andrzej Sokołowski

**Funding Acquisition**: Tristan Glatard, Jean-Baptiste Poline, Madeleine Sharp

**Investigation**: Andrzej Sokołowski, Jean-Baptiste Poline, Madeleine Sharp, Tristan Glatard **Methodology**: Andrzej Sokołowski, Tristan Glatard

**Project Administration**: Tristan Glatard

**Resources**: Tristan Glatard

**Software**: Andrzej Sokołowski, Mathieu Dugré, Yohan Chatelain, Tristan Glatard

**Supervision**: Tristan Glatard

**Validation**: Andrzej Sokołowski, Tristan Glatard

**Visualization**: Andrzej Sokołowski

**Writing - Original Draft Preparation**: Andrzej Sokołowski

**Writing - Review** & **Editing**: Andrzej Sokołowski, Nikhil Bhagwat, Yohan Chatelain, Mathieu Dugré, Alexandru Hanganu, Oury Monchi, Brent McPherson, Michelle Wang, Jean-Baptiste Poline, Madeleine Sharp, Tristan Glatard

